# Pioneer factor Foxa2 enables ligand-dependent activation of LXRα

**DOI:** 10.1101/2020.04.10.036061

**Authors:** Jessica Kain, Xiaolong Wei, Andrew J. Price, Claire Woods, Irina M. Bochkis

**Author notes:** To whom correspondence should be addressed: Irina M. Bochkis, Ph. D. Department of Pharmacology, University of Virginia School of Medicine, 5030 Pinn Hall, 1340 Jefferson Park Ave, Charlottesville, VA 22908.

## Abstract

Type II nuclear hormone receptors, such as FXR, LXR, and PPAR, which function in glucose and lipid metabolism and serve as drug targets for metabolic diseases, are permanently positioned in the nucleus regardless of the ligand status. Ligand activation of these receptors is thought to occur by co-repressor/co-activator exchange, followed by initiation of transcription. However, recent genome-wide location analysis showed that LXRα and PPARα binding in the liver is largely ligand-dependent. We hypothesized that pioneer factor Foxa2 evicts nucleosomes to enable ligand-dependent receptor binding. We show that chromatin accessibility, LXRα occupancy, and LXRα-dependent gene expression upon ligand activation require Foxa2. Unexpectedly, Foxa2 occupancy is drastically increased when LXRα is bound by an agonist. Our results suggest that Foxa2 and LXRα bind DNA as an interdependent complex during ligand activation. Our model requiring pioneering activity for ligand activation challenges the existing co-factor exchange mechanism and expands current understanding of nuclear receptor biology, suggesting that chromatin accessibility needs to be considered in design of drugs targeting nuclear receptors.

## Introduction

Members of nuclear receptor superfamily, farnesoid X receptors (FXR) liver X receptors (LXR) and peroxisome proliferator-activated receptors (PPAR) function in bile acid, fatty acid, cholesterol, and glucose metabolism (*1*). Ligands that activate or antagonize these receptors have been developed to treat metabolic disease and cancer (*2–5*). Nuclear receptors are classified according to their subcellular localization in the absence of ligand and mechanism of action. Type I receptors, including estrogen (ER) and androgen (AR) receptors, are positioned in the cytoplasm and bound to chaperone heat-shock proteins, translocating to the nucleus and binding hormone-response DNA elements as homodimers upon ligand binding. In contrast, type II receptors, such as FXR, LXR, and PPAR, are permanently positioned in the nucleus regardless of the ligand status. They bind DNA as heterodimers in complex with retinoid X receptor (RXR). The accepted paradigm regarding ligand activation of type II receptors is a two-step process: 1) the receptor is bound to DNA in complex with a co-repressor in absence of the ligand; 2) binding of the ligand induces a conformational change, co-repressor/co-activator exchange, and initiation of transcription (**Fig. 1A**, top panel). However, recent genome-wide location analysis showed that, like for type I receptors ER and AR (*6*) (*7*), LXRα and PPARα binding in the liver is largely ligand-dependent (*8*). While binding of ER and AR in ligand-dependent manner is compatible with their mechanism of action (translocation from the cytoplasm to the nucleus and DNA binding upon ligand binding), a similar process governing recruitment of type II receptors, which are permanently nuclear, is unexplained.

We hypothesized that pioneer factor Foxa2 modulates chromatin accessibility by evicting nucleosomes to enable binding by type II nuclear receptors upon ligand activation. Foxa2 is a member of the Foxa subfamily of winged-helix/*forkhead* box (Fox) transcription factors (*9*), named “pioneer” factors for their ability to independently bind highly condensed chromatin, displacing linker histones, and facilitate access for subsequent binding of additional transcription factors (*10*). Foxa2 binds nucleosomal DNA *in vivo* (*11*) and enables nucleosomal depletion during differentiation (*12*). We have previously demonstrated that ligand-responsive activation of FXR gene expression is Foxa2-dependent (*13*) and that Foxa2 cooperates with ligand-activated PPARα receptors (*14*) in aged liver. Here we show that chromatin accessibility, recruitment of LXRα to genomic sites, and LXRα-dependent gene expression upon ligand binding (using two ligands, GW3965 and T0901317) require Foxa2. Unexpectedly, we observe that Foxa2 occupancy is drastically increased when LXRα is activated by an agonist. Considering that Foxa2 can open closed chromatin and the agonist does not interact with Foxa2 but binds the ligand-binding domain of LXRα, our results suggest that Foxa2 and LXRα bind DNA as an interdependent complex during ligand activation. Our model requiring pioneering activity for ligand activation of type II nuclear receptors challenges the existing co-factor exchange mechanism and expands current understanding of nuclear receptor biology. Our results suggest that the role of pioneer factors in changing chromatin accessibility needs to be considered in design of drugs that target nuclear receptors.

**Figure 1.**
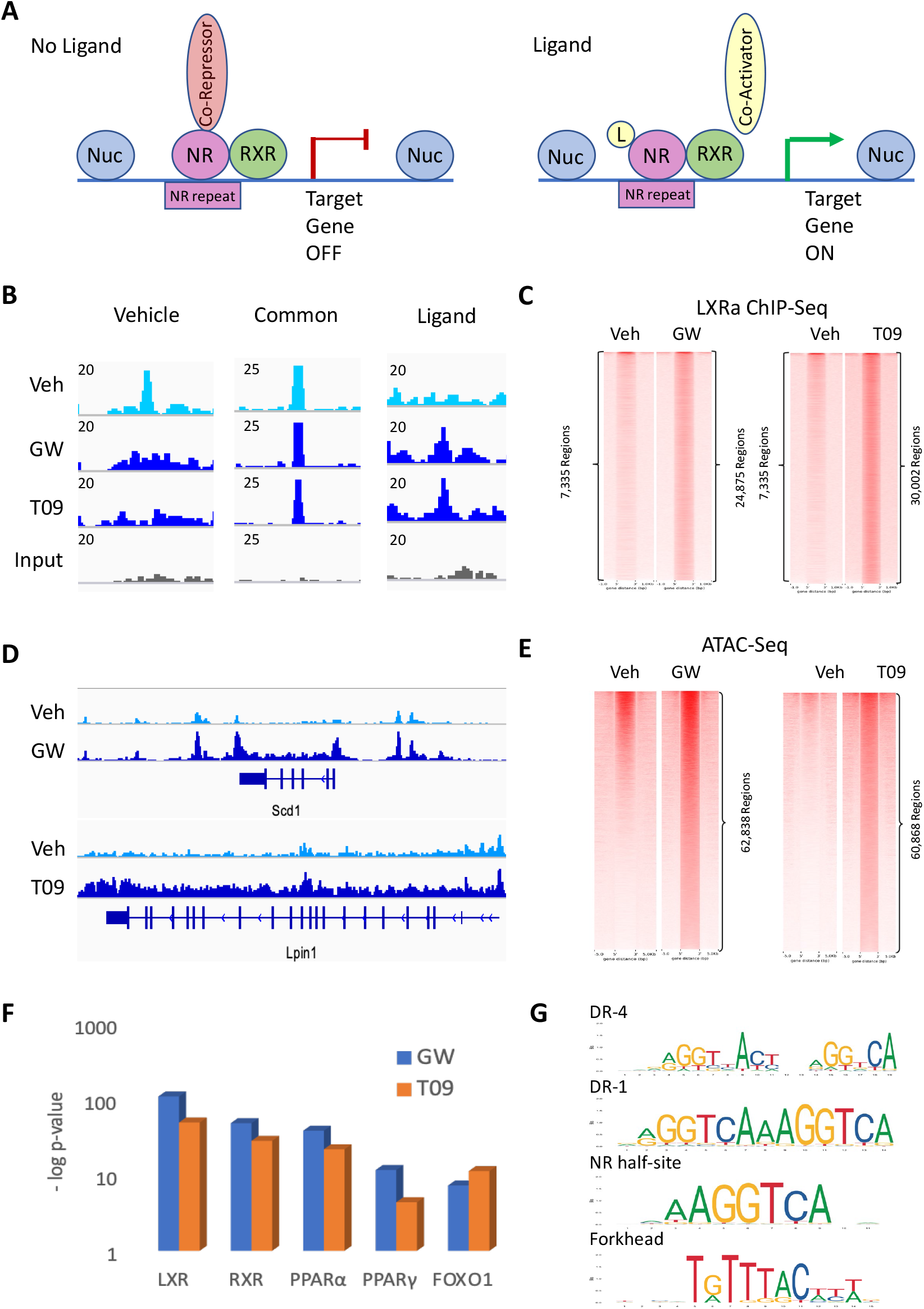
Chromatin accessibility increases with addition of LXR ligands. (**A**) The accepted paradigm regarding ligand activation of type II nuclear receptors (NR) is a two-step process: 1) the receptor as a heterodimer with retinoid X receptor (RXR) is bound to the DNA in complex with a co-repressor in absence of the ligand; 2) binding of the ligand induces a conformational change, co-repressor/co-activator exchange and initiation of transcription. (**B**) Examples of 3 types of sites bound by LXRα: 1) only in control mice treated with vehicle (left panel chr8:47,640,758-47,683,677), 2) common (chr8:46,531,348-46,574,267 middle panel), and 3) ligand-dependent or occupied only during ligand activation (chr2:45,871,831-45,881,682 right panel), ChiP-Seq track view in Integrative Genome Viewer (IGV). Signal was merged from four biological replicates in each condition. Track of input reads is provided for comparison. (**C**) Heatmaps showing increased LXRα binding with ligand activation (GW, 24,875 regions, left panel;) T09, 30,002 regions, right panel). ChIP-Seq peaks called by PeakSeq (FDR < 5%, q-value < 0.005, vs input control). (**D**) ATAC-seq track view in IGV of induced chromatin accessibility with ligand activation (GW, Scd1 locus, top panel; T09, Lpin1, bottom panel). (**E**) Heatmaps showing genome-wide ATAC-Seq signal with ligand activation by GW (62,838 regions, left panel) and T09 (60,868 regions, left panel; (PeakSeq, FDR < 5%, q-value < 0.05, vs input control). Signal was merged from two biological replicates in each condition. (**F**) Analysis of open chromatin regions induced by LXR ligands with published ChIP-Seq data sets in the mouse genome identified binding sites of nuclear receptors LXR, RXR, PPARα, PPARγ, and *forkhead* factor FOXO1 as significantly enriched. (graph of −log p-values, GW, blue; T09, orange; ChEA analysis in enrichR). (**G**) Scanning motif of positional weight matrices in Jaspar and TRANSFAC databases by PscanChIP identified consensus sites for LXR (DR-4 element, p-value < 1.9E-84; half-site, p-value <6.3E-211) and other nuclear receptors (DR-1 element, p-value <1.2E-143) as well as *forkhead* motifs (p-value < 3.8E-213).

## Results

### Chromatin accessibility increases with addition of LXR ligands

A recent genome-wide binding study demonstrated that LXRα occupancy in the liver is largely ligand-dependent (*8*), disputing the accepted co-factor exchange mechanism for ligand activation of type II nuclear receptors (**Fig. 1A**) (*1*). We hypothesized that increase in chromatin accessibility leads to additional LXRα binding during ligand activation. First, we performed LXRα genome-wide location analysis to assess acute changes in LXRα occupancy in livers of mice treated with the agonist for four hours (either GW3965, specific for LXR, or T0901317, which binds LXR and few other related receptors). We found 3 types of sites bound by LXRα: 1) only in control mice treated with vehicle (left panel), 2) common (middle panel). and 3) ligand-dependent or occupied only during ligand activation (right panel, **Fig. 1B**). Overall, we observed that LXRα binding greatly increases during ligand activation (7, 335 regions for vehicle, 24,875 regions for GW, 30,002 regions for T09; PeakSeq, FDR < 5%, q-value < 0.005, **Fig. 1C**). Our results agree with the previous study where authors assessed LXRα occupancy after 14 days of daily ligand injections (*8*). Next, we profiled chromatin accessibility in livers of mice treated with LXR ligands (ATAC-Seq), demonstrating that addition of the agonist leads to significant chromatin opening at loci of known LXR targets Lpin1 and Scd1 (**Fig. 1D**) and in a genome-wide manner (62,838 regions for GW compared to vehicle, 60,868 regions for compared to vehicle, PeakSeq, FDR < 5%, q-value < 0.05, **Fig. 1E**). The overlap analysis of open chromatin regions induced by LXR ligands with published ChIP-Seq data sets in the mouse genome identified binding sites of nuclear receptors LXR, RXR, PPARα, PPARγ, and *forkhead* factor FOXO1 as significantly enriched (**Fig. 1F**, ChEA analysis in enrichR (*15*)). Our results are consistent with the previous results showing that LXR binding sites are extensively shared with other nuclear receptors in the liver (*8*). In addition, genes in regions opened during LXR ligand activation substantially overlap with genes bound by ENCODE transcription factors in human genome (FOXA2 in HepG2 cells, p-value < 4.5E-4, HNF4α in HepG2 cells, p-value < 0.01, and LXR target SREBF1 in HepG2 cells, p-value <0.01). Scanning motif of positional weight matrices in Jaspar and TRANSFAC databases by PscanChIP (**Fig 1G**) (*16*) identified consensus sites for LXR (DR-4 element, p-value < 1.9E-84; half-site, p-value <6.3E-211) and other nuclear receptors (DR-1 element, p-value <1.2E-143) as well as *forkhead* motifs (p-value < 3.8E-213). Together, these analyses confirm that newly open chromatin sites are bound by LXR and suggest a role for forkhead factors in chromatin opening during ligand activation.

### Foxa2 opens chromatin for LXRα binding during ligand activation

We have previously determined that ligand-responsive activation of FXR gene expression is Foxa2-dependent (*13*) and that Foxa2 cooperates with ligand-activated PPARα receptors (*14*) in aged liver. We hypothesized that pioneer factor Foxa2 modulates chromatin accessibility by evicting nucleosomes to enable binding by type II nuclear receptors upon ligand activation. Hence, we performed ATAC-Seq in livers of *Foxa2* mutants and their control littermates treated with LXR ligands (*Foxa2^loxP/loxP^;Alfp.Cre* described in (*17*)), showing that changes in chromatin accessibility at the locus of known LXR targets *Scd1* (**Fig. 2A**, left panel) and *Lpin1* (data not shown), as well as in a genome-wide manner (**Fig. 2B**), are absent in *Foxa2*-deficient livers. Next, we performed genome-wide location analysis of Foxa2 in wildtype mice ((*Foxa2^loxP/loxP^*) treated with LXR agonists and observed that additional Foxa2 binding sites at the *Scd1* locus in livers treated with GW corresponds to increase in chromatin accessibility that is absent in *Foxa2* mutants (**Fig. 2A**, right panel).

**Figure 2.**
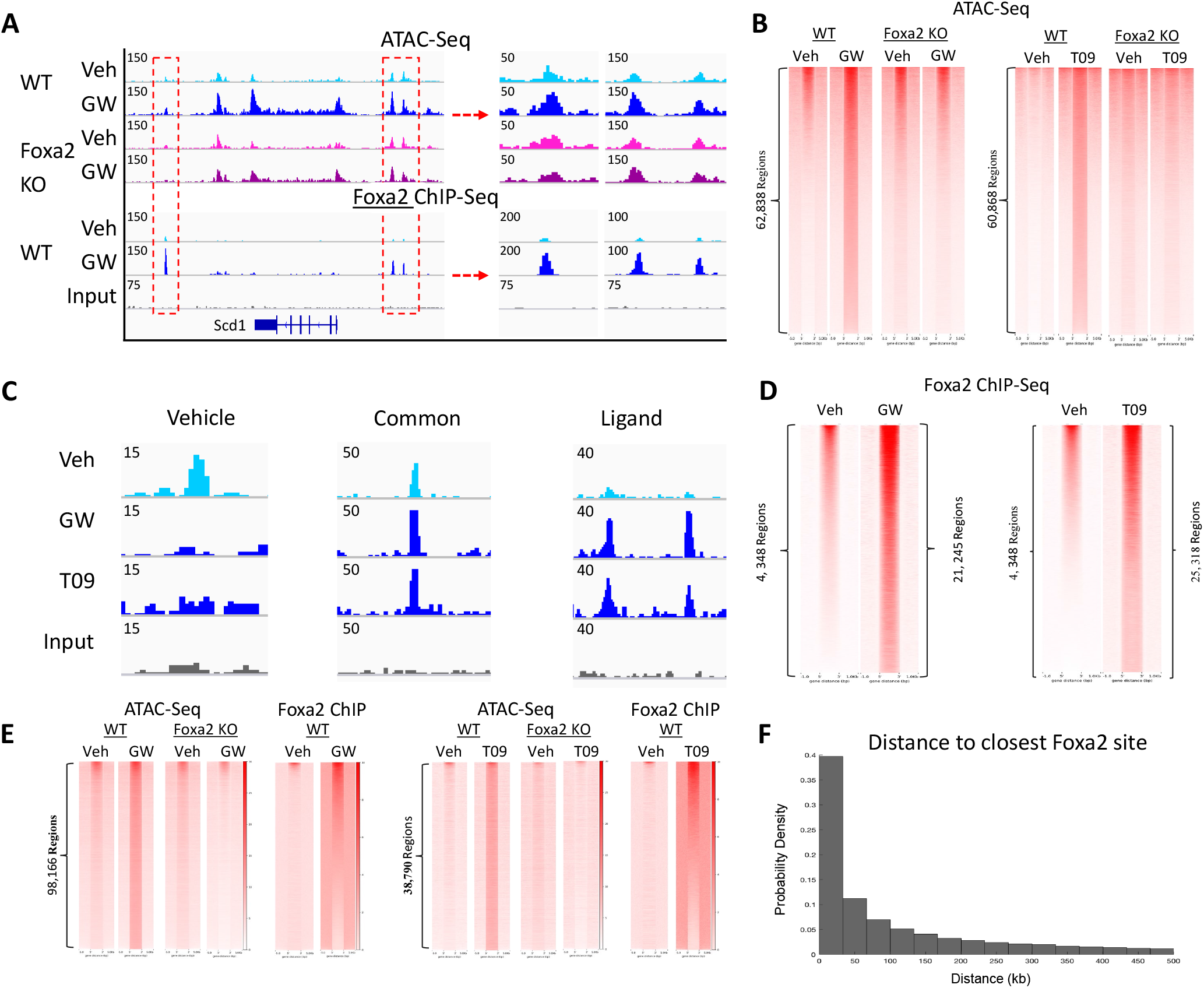
Increase in chromatin accessibility with addition of LXR ligand requires Foxa2. (**A**) Increase in chromatin accessibility upon ligand activation observed at the Scd1 locus featured in **Figure 1D** (IGV view of ATAC-Seq signal, top two tracks, chr19:44,374,231-44,427,408) is absent in Foxa2 mutants (middle two tracks). Ligand-dependent increase in chromatin accessibility correlates to additional Foxa2 binding during GW treatment (IGV view of Foxa2 ChIP-Seq signal, bottom tracks, track of input reads is provided for comparison.). Middle and right panel (chr19:44,376,869-44,383,515; chr19:44,414,227-44,420,873) are zoomed in regions marked by red rectangles on the left panel. (**B**) Heatmaps showing genome-wide increase in ATAC-Seq signal with LXRα ligand activation by GW (right panel, 62,838 regions) and T09 (left panel, 60,868 regions) in wildtype controls is absent in *Foxa2*-deficient livers. (**C**) Examples of 3 types of sites bound by Foxa2: 1) only in control mice treated with vehicle (left panel chr1:72,461,246-72,463,888), 2) common (chr4:82,395,078-82,399,310 middle panel), and 3) ligand-dependent or occupied only during ligand activation (chr4:6,273,103-6,281,569 right panel), ChIP-Seq track view in Integrative Genome Viewer (IGV). Signal was merged from three biological replicates in each condition. Track of input reads is provided for comparison. (**D**) Heatmaps showing Foxa2 binding is induced by LXR agonists (GW, 21,245 regions, left panel; T09, 25,318 regions, right panel; (PeakSeq, FDR < 5%, q-value < 0.0005; wildtype GW vs Input control; wildtype T09 vs Input control). (**E**) Heatmaps showing co-localization of Foxa2 ChIP-Seq and ATAC-Seq signal in regions with Foxa2-dependent chromatin accessibility during ligand activation (ATAC-Se regions, GW, 98,166 regions, left panel; T09, 38,790, left panel; PeakSeq, FDR < 5%, q-value < 0.05; WT GW vs Foxa2 GW; WT T09 vs Foxa2 KO T09). (**F**) Histogram showing relative frequency of the distances from the 98,166 ATAC-Seq peak regions (GW activation, **Figure 2E**) to the closest Foxa2 binding site (called by PeakSeq, q-value < 0.0005).

Similar to LXRα, we found 3 types of sites bound by Foxa2: 1) only in control mice treated with vehicle (left panel), 2) ligand-dependent or occupied only during ligand activation (right panel), and 3) common to both conditions (middle panel, **Fig. 2C**). Unexpectedly, Foxa2 occupancy dramatically increased with addition of either ligand (4, 348 regions for vehicle, 21,245 regions for GW, 25,318 regions for T09; PeakSeq, FDR < 5%, q-value < 0.0005, **Fig. 2D**). Next, we proceeded to ascertain the overlap of open regions induced by LXR ligands in controls and absent in *Foxa2* mutants with additional Foxa2 binding (**Fig. 2E**), computing the intersection using Foxa2 ChIP-Seq coverage at ATAC-Seq regions. We found that 22,752 and 16,214 regions with induced chromatin accessibility (GW and T09, respectively, cutoffs 2 reads/bp) were bound by Foxa2. The number of Foxa2 occupied regions in the overlap is comparable or exceeds the number of Foxa2 sites we called bound during ligand activation by PeakSeq at our chosen cutoff (FDR < 5%, q-value < 0.0005, **Fig. 2D**). Since the number of accessible sites (98,166 ATAC-Seq regions comparing WT GW to KO GW, 38,790 regions for WT T09 to KO T09 comparison, PeakSeq, FDR < 5%, q-value < 0.05) far exceeds additional Foxa2 occupancy during ligand activation, we plotted the distribution of open chromatin regions to the nearest Foxa2 site. Although Foxa2 binds at enhancers (*18*) and is known to act distally (*19*), over 40% of accessible sites are proximal, within 35 Kb of the Foxa2 site (**Fig. 2F**).

We observe a dramatic change in Foxa2 occupancy in livers treated with LXR agonists (**Fig. 2D**). Foxa2 is not known to bind nuclear receptor agonists directly and has no ligand-binding domain. Considering that Foxa2 opens closed chromatin and the agonist does not interact with Foxa2 but binds the ligand-binding domain of LXRα, we hypothesized that Foxa2 and LXRα binding to DNA is interdependent during ligand activation.

### Foxa2 interacts with LXRα in ligand-dependent manner

To test whether Foxa2 and LXRα formed a complex that bound DNA upon agonist addition (**Fig. 3A**), we performed reciprocal immunoprecipitation experiments and determined that Foxa2 and LXRα interact in a ligand-dependent manner. First, liver extracts were immunoprecipitated with an antibody to LXRα and subsequent immunoprecipitate was blotted with an antibody to Foxa2, showing a strong interaction between LXRα and Foxa2 in both GW and T09 treatment but not in IgG and vehicle controls (**Fig. 3B**, left panel). Then, we reversed the order, immunoprecipitating with antibody to Foxa2 and blotting for LXRα, exhibiting a weaker interaction between Foxa2 and LXRα also in ligand-dependent manner (**Fig. 3B**, right panel). Our results suggest that Foxa2 is required for LXRα binding, with majority of hepatic liganded LXRα in complex with Foxa2. In contrast, a smaller proportion of hepatic Foxa2 is bound to LXRα, as the pioneer factor performs additional functions in the cell.

**Figure 3.**
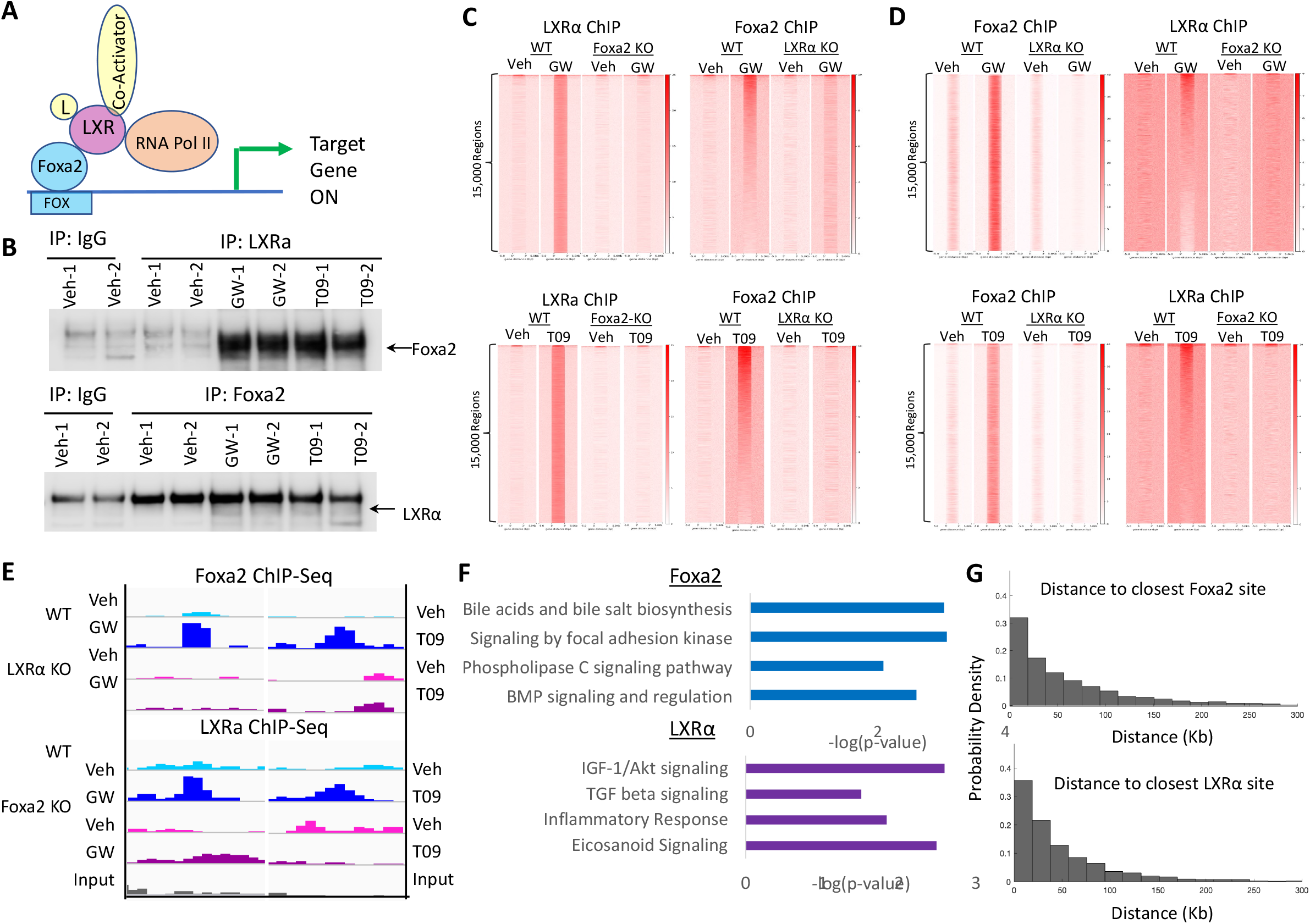
Foxa2 interacts with LXRα in ligand-dependent manner. (**A**) Foxa2/LXR interaction model. Foxa2 and liganded LXRα form a complex, with pioneer factor binding DNA at the *forkhead* (Fox) binding site. (**B**) Reciprocal co-immunoprecipitation experiments in livers of mice treated with LXR ligands. Immunoprecipitation with an antibody to LXRα and subsequent blotting with an antibody to Foxa2, shows a strong interaction between LXRα and Foxa2 in both GW and T09 treatment but not in IgG and vehicle controls (left panel). Then, in reverse the order, immunoprecipitation with antibody to Foxa2 and blotting for LXRα, exhibits a weaker interaction between Foxa2 and LXRα also in ligand-dependent manner (right panel). (**C**) Heatmaps of ChIP-Seq signal at top 15,000 LXRα-binding regions in wildtype controls (left panel) that is lost in *Foxa2* mutants (left panel) and overlap of Foxa2 ChIP signal in wildtype controls at these sites that is lost in LXRα mutants (right panel). (**D**) Heatmaps of ChIP-Seq signal at top 15,000 Foxa2-binding regions in wildtype controls (left panel) that is lost in LXRα mutants (left panel) and overlap of LXRα ChIP signal in wildtype controls at these sites that is lost in *Foxa2* mutants (right panel). (**E**) Examples of co-localized Foxa2/LXR ChIP signal in wildtype controls present only during ligand activation and absent in reciprocal knockouts (Foxa2 ChIP in LXRα mutants & LXRα ChIP in *Foxa2* mutants, chr1:12,643,509-12,646,151; chr16:80165264-80165280), suggesting that Foxa2 and LXRα binding to DNA is interdependent during ligand activation. Signal was merged for three (Foxa2) or four (LXR) biological replicates). Track of input reads is provided for comparison. (**F**) The pathways associated with genes mapped to LXR-only or Foxa2-only regions (LXRα signal below 1 read/bp for GW and below 2 reads/bp for T09). Genes regulating bile acid homeostasis were associated with Foxa2-bound regions, consistent with our previous studies (*13, 20*). (**G**) Histogram showing relative frequency of the distances from “LXR-only” sites from that do not overlap with Foxa2 signal (GW activation, **Fig. 3D**) to the closest Foxa2 binding site (called by PeakSeq, q-value < 0.0005, top panel). Histogram showing relative frequency of the distances from “Foxa2-only” sites from that do not overlap with LXRα signal (GW activation, **Figure 3E**) to the closest LXRα binding site (called by PeakSeq, q-value < 0.005, bottom panel).

Next, we ascertained common genomic binding of Foxa2 and LXRα, also in reciprocal fashion. We looked for Foxa2 signal at top 15,000 LXRα binding sites induced by LXR agonists in wildtype controls (**Fig. 3C**) and LXRα signal at top 15,000 Foxa2 binding sites induced by LXR ligands (**Fig. 3D**). The overlap was more prominent for Foxa2 binding at LXRα sites as compared to LXRα occupancy at Foxa2 sites, consistent with the results of the co-immunoprecipitation experiments (**Fig. 3B**). While we observed Foxa2 binding in LXRα mutants and LXRα binding in *Foxa2*-deficient mice (**fig. S1**), binding by either factor was absent in reciprocal knockout animals at Foxa2 & LXRα binding sites occupied in wildtype controls. In addition, binding of Foxa2 was severely affected by lack of LXRα (PeakSeq, FDR < 5%, q-value < 0.0005, 2,009 regions vehicle, 58 regions for GW, and 1,695 for T09, all in LXRα KO mice vs. Input control) and LXRα occupancy was substantially reduced without Foxa2 (PeakSeq, FDR < 5%, q-value < 0.005, 5,354 regions vehicle, 1,024 regions for GW, and 11,402 for T09, all in Foxa2 KO mice vs.\Input control).

Using ChIP-Seq coverage, we separated Foxa2 and LXRα targets into Foxa2/LXRα dual sites, LXR-only sites (Foxa2 signal below 1.5 read/bp for GW and below 2 reads/bp for T09 in LXRα-bound regions in **Fig. 3C**) and Foxa2-only regions (LXRα signal below 1 read/bp for GW and below 2 reads/bp for T09 in Foxa2-bound regions in **Fig. 3D**). Example of a dual Foxa2/LXRα binding sites occupied by both factors in wildtype controls but not in reciprocal mutants are shown **Fig. 3E** (GW left panel, T09 right panel). Pathways associated with genes mapped to LXR-only or Foxa2-only regions are shown in **Fig. 3F**. Genes regulating bile acid homeostasis were associated with Foxa2-bound regions, consistent with our previous studies (*13, 20*). For LXR-only sites, we computed the distance to the closest Foxa2 binding site (called by PeakSeq, FDR < 5%, q-value < 0.0005, **Fig. 2D**) and found over a third (34%) within 20 Kb (**Fig. 3G**, top panel). For Foxa2-only sites, about a quarter (27%) had an LXR peak (PeakSeq, FDR < 5%, q-value < 0.005, **Fig. 1C**) within 20 Kb (**Fig. 3G**, bottom panel).

PscanChIP analysis demonstrated that of all scanned positional weight matrices in Jaspar database, the forkhead motifs were most significantly enriched in both dual and Foxa2-only regions (**fig. S2**). Nuclear receptor consensus sites were highly enriched in dual sites (HNF4G, p-value < 2.3 E-172, NR1H3:RXR, p-value <1.3 E-25) and LXR-only regions (RORA, p-value < 1.3 E-200). Unexpectedly, forkhead motifs are also prevalent in LXR-only sites (FOXA1, p-value <9.7 E-265, FOXA2, p-value <1.6 E-224). Interestingly, the overlap analysis of both Foxa2-only and LXR-only regions with published ChIP-Seq data sets in the mouse genome identified binding sites of sane factors (Smarca4, Cebpd, Stat3, Zfp57, ChEA analysis in enrichR). Foxa2 has been shown to partner with chromatin remodelers like Smarca4 (BRG-1) that co-activate LXRα-dependent gene expression (*12, 21*). It is likely that with multiple co-factors in Foxa2/LXRα complex, the interaction between Foxa2 and LXRα at all dually bound sites cannot be captured by single factor ChIP-Seq. The other possibility is that Foxa2 and LXRα interact distally in a ligand-dependent manner, as suggested by distance analysis in **Fig. 3G**, and, therefore, their binding sites map to different genomic locations.

### Ligand-dependent activation of LXR gene expression is Foxa2-dependent

To test whether changes in chromatin accessibility and transcription factor binding have functional consequences, we performed RNA-Seq analysis in livers of *Foxa2* mutants and their control littermates treated with LXR agonists. First, we assessed the expression of LXRα targets *Scd1* and *Lpin1* since increase in chromatin accessibility induced at these loci during ligand activation was absent in *Foxa2*-deficient livers and newly opened chromatin regions corresponded to additional, ligand-dependent Foxa2 binding (**Fig. 2A**). Indeed, while mRNA levels of both genes are increased by ligand treatment in wildtype controls, the increase is reduced in *Foxa2* mutants (**Fig. 4A**). In addition, expression of numerous Foxa2-dependent LXRα target genes (induced by ligand in wildtype mice but completely blunted in *Foxa2* mutants) activated either by GW (left panel) or T09 (right panel) is shown in heatmaps in **Fig. 4B**. While GW induced expression of only 104 genes in wildtype mice, expression of 308 genes were changed in *Foxa2* mutants treated with this compound (**Fig. 4C**, top panel). Overwhelming majority (267/308) were differentially expressed only in *Foxa2*-deficient mice. We observed a similar pattern in mice treated with T09. Overall, mRNA levels of 427 genes were differentially expressed in wildtype mice treated with T09, but only 195 showed comparable changes in *Foxa2* mutants (**Fig. 4C**, bottom panel). Also, deletion of *Foxa2* resulted in additional 217 genes that were not changed in controls but differentially regulated in *Foxa2* mutants treated with T09 (**Fig. 4C**, bottom panel). Hence, Foxa2 deficiency affects ligand-dependent activation of LXRα gene expression in two ways. First, expression of hundreds of genes induced by the ligand in wildtype controls is blunted in *Foxa2* mutants (**Fig. 4B**). And, second, mRNA levels of numerous genes that are not changed by the ligands in control mice are changed in *Foxa2*-deficient livers (**Fig. 4C**).

**Figure 4.**
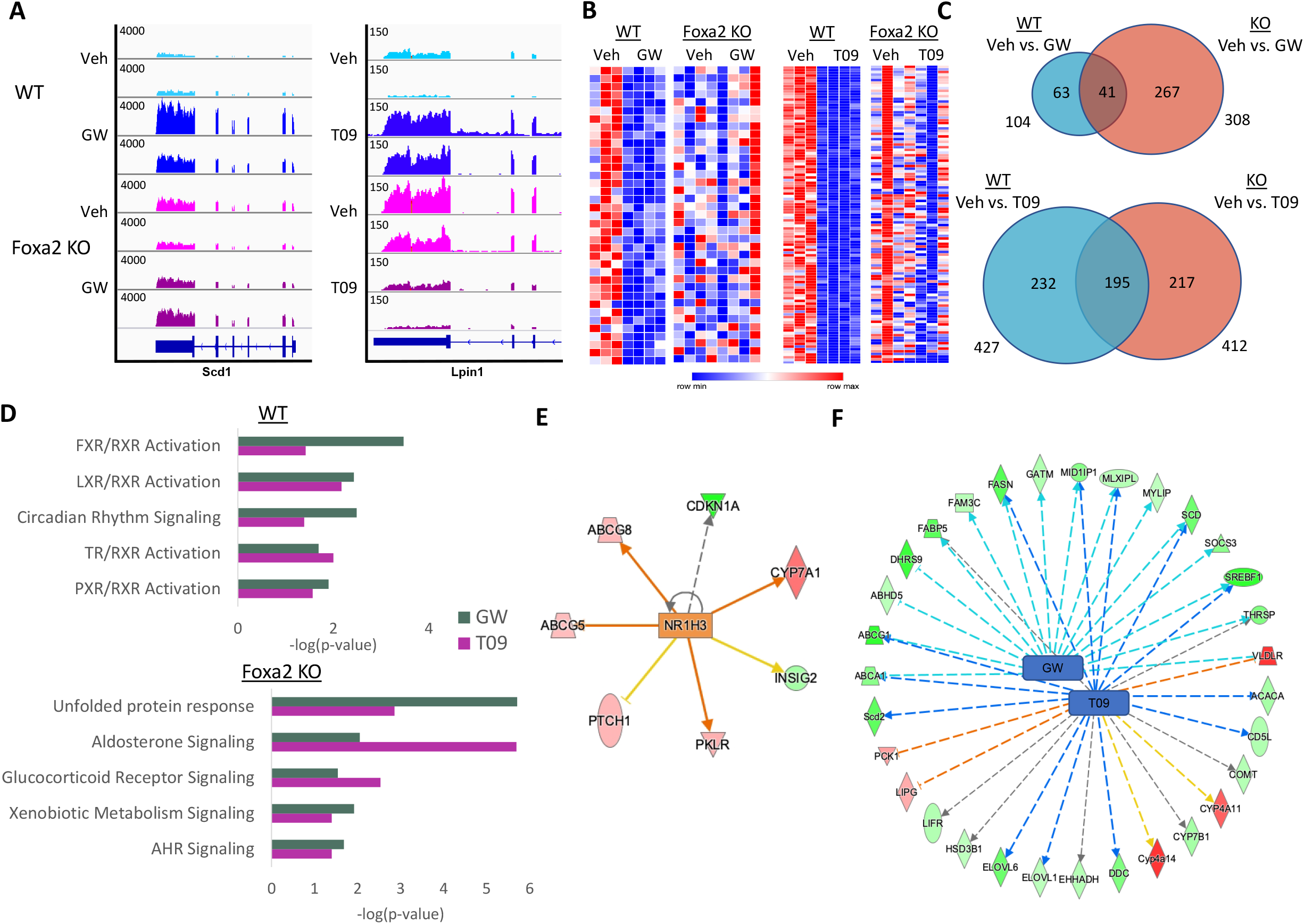
Ligand-dependent activation of LXRα gene expression is Foxa2-dependent. (**A**) RNA-seq track view in IGV showing that ligand-responsive activation of Scd1 (GW, left panel) and Lpin1 (T09, right panel), well-known LXRα targets, is Foxa2-dependent. (**B**) Heatmap (RNA-Seq gene expression) of Foxa2-dependent ligand-activated LXRα targets. Expression of genes induced by LXR ligand (GW, 37 genes, left panel; T09, 111 genes, right panel; EdgeR, FDR<5%) in wildtype mice but not upregulated in Foxa2 mutants.(**C**) Venn diagram showing the number of differentially expressed genes with ligand activation by GW (top panel, 104 genes in WT, 308 genes in KO, 41 genes in common) and T09 (bottom, 427 genes in WT, 412 genes in KO, 195 genes in common). Differential expression was analyzed using EdgeR (FDR 5%). (**D**) Ingenuity Pathway Analysis of genes that mapped to differentially expressed genes in wildtype (top) and Foxa2-knockout (bottom) mice identified activation of nuclear receptors pathways, including LXR, FXR, PXR, and TR (top panel; p-values <0.0037 and 0.0068, 0.00034 and 0.038, 0.013 and 0.0227, 0.02 and 0.01 for GW and T09, respectively. In contrast, “Unfolded Protein Response”, “Aldosterone Signaling”, and “Glucocorticoid Signaling” were among enriched pathways in *Foxa2* mutants treated with GW and T09 (bottom panel; p-values < 1.95 E-6 and 0.0014,, 0.0089 and 2.04 E-6, 0.029 and 0.003, for GW and T09 respectively). (**E**) IPA regulatory network analysis of differentially expressed genes during GW activation identified LXR/RXR heterodimer as an activator of ligand-dependent gene expression in wildtype controls (blue line represents activation, orange line repression, and gray line the association with expression change). (**F**) IPA network analysis identified GW and T09-dependent gene expression (31 LXR targets) to be inhibited in *Foxa2*-defcient mice. (blue line represents activation, orange line repression, and gray line the association with expression change).

Ingenuity Pathway Analysis (IPA) of differentially expressed transcripts in wildtype livers treated with LXR ligands identified activation of nuclear receptors pathways, including LXR, FXR, PXR, and TR (p-values <0.0037 and 0.0068, 0.00034 and 0.038, 0.013 and 0.0227, 0.02 and 0.01 for GW and T09, respectively, **Fig. 4D**, top panel). In contrast, “Unfolded Protein Response”, “Aldosterone Signaling”, and “Glucocorticoid Signaling” were among enriched pathways in *Foxa2* mutants treated with GW and T09 (p-values < 1.95 E-6 and 0.0014,, 0.0089 and 2.04 E-6, 0.029 and 0.003, for GW and T09 respectively, **Fig. 4D**, bottom panel) Next, we employed computational network analysis to investigate which transcription factors mediated the gene expression changes observed in mice treated with LXR agonists. IPA network analysis identified LXR/RXR heterodimer activating ligand-dependent gene expression in wildtype controls (**Fig. 4E**) and GW and T09-dependent gene expression (31 LXR targets) to be inhibited in *Foxa2*-deficient mice treated with these ligands (**Fig. 4F**). In summary, changes in chromatin accessibility and transcription factor binding observed with addition of LXR agonists lead to activation of gene expression in wildtype controls that is inhibited in *Foxa2* mutants.

### Paradigm-shifting model requiring pioneering activity for ligand-dependent activation of LXRα

In summary, our data show that in addition to ligand-independent sites (**Fig. 5A**, top left panel, IGV example on the right), LXRα binds to ligand-dependent nuclear receptor (NR) sites that are inaccessible without ligand activation and become exposed upon stimulation. Pioneer factor Foxa2 enables ligand-dependent NR binding, evicting the nucleosome that occludes NR binding site upon ligand activation. Two mechanisms are plausible: I) Foxa2 is bound in absence of ligand and evicts the nearby nucleosome occluding the nuclear receptor response element upon ligand binding (**Fig. 5A**, middle left panel, IGV example on the right); II) Foxa2 is not bound prior to ligand activation ((**Fig. 5**, bottom panel, IGV example on the right). Dramatic change in Foxa2 occupancy observed in livers treated with LXR agonists (**Fig. 2C**) provides overwhelming evidence for second mechanism.

**Figure 5.**
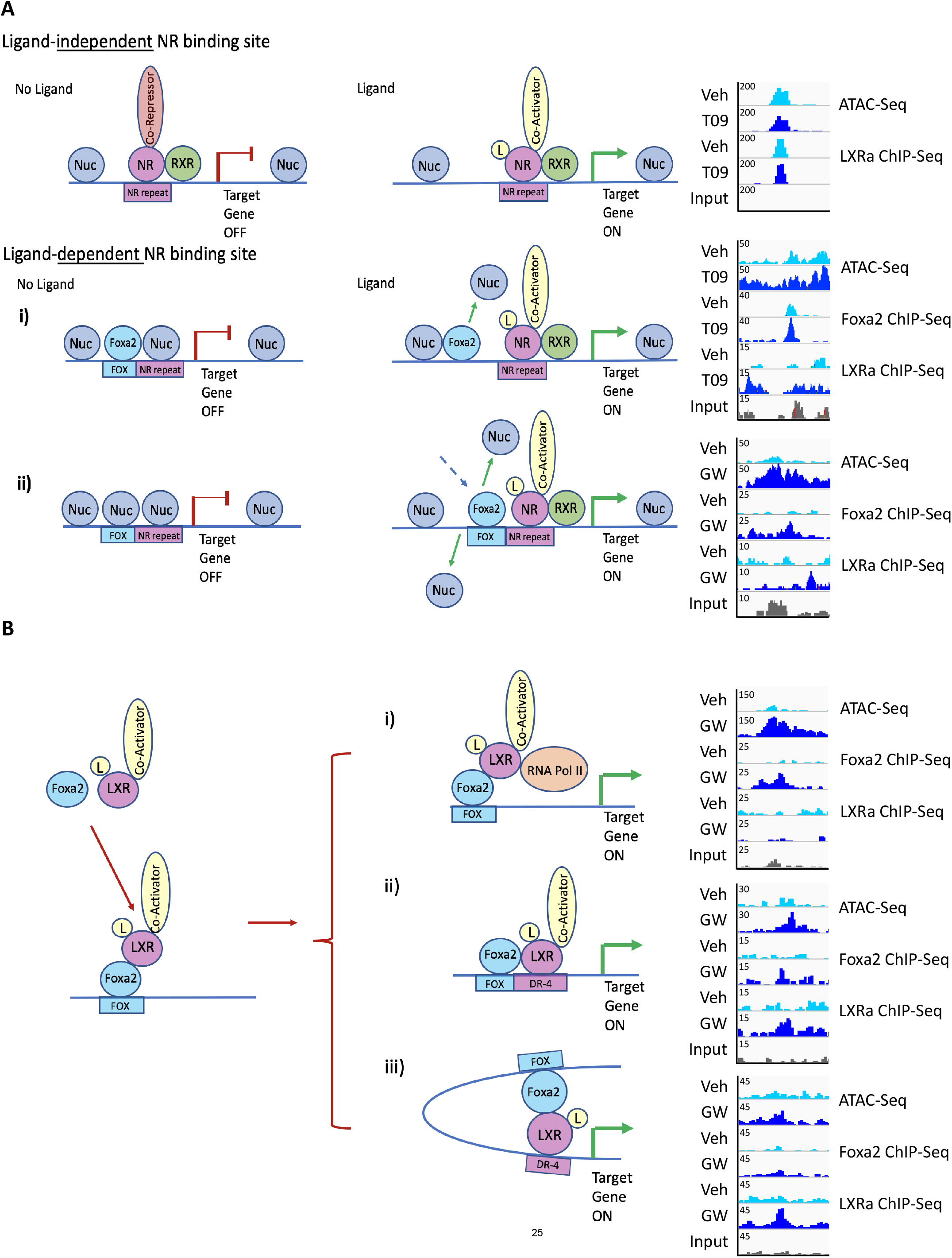
Model describing how Foxa2 enables ligand-dependent activation of LXRα. (**A**) We distinguish between ligand-independent and ligand-dependent nuclear receptor (NR) binding sites. Ligand-independent NR sites are always accessible and the mechanism of LXR activation is consistent with the classical model (top panel). In the absence of ligand, the receptor is bound to the DNA in complex with a co-repressor (top panel, left), while ligand binding induces a conformational change, co-repressor/co-activator exchange, and initiation of transcription (top panel, middle). An example (top panel, right) shows a region where both chromatin accessibility and LXRα binding do not change upon ligand addition. In contrast, ligand-dependent NR sites are inaccessible without ligand activation and become exposed upon stimulation (middle and bottom panel). We demonstrate that pioneer factor Foxa2 enables ligand-dependent NR binding, evicting the nucleosome (Nuc) that occludes NR binding site upon ligand activation. Two mechanisms are plausible: i) Foxa2 is bound in absence of ligand and evicts the nearby nucleosome occluding NR response element upon ligand binding (middle panel showing a pre-existing Foxa2 binding site, increase in chromatin accessibility and a new LXRα binding site upon agonist addition); ii) Foxa2 is not bound prior to ligand activation.(bottom panel showing a novel ligand-dependent Foxa2 site corresponding to increase in chromatin accessibility and additional LXRα binding during ligand activation). (**B**) Upon ligand binding, Foxa2 forms a complex LXRα with, binds previously inaccessible DNA, enabling LXRα to access its response element. Our computational analysis provides evidence for 3 modes of synergistic regulation by Foxa2 and LXRα during ligand activation: Foxa2 initially bound DNA in complex with liganded LXRα, acting as a co-activator. LXRα either remains in the same position in the complex (i, top panel, IGV example on the right showing ligand-dependent Foxa2-only site that corresponds to increase in chromatin accessibility) or moves to an adjacent (ii, middle panel, IGV example demonstrating ligand-dependent dual Foxa2/LXRα binding site that corresponds to increase in chromatin accessibility) or a distal binding site (iii, bottom panel, IGV example demonstrating ligand-dependent LXRα-only binding site that corresponds to increase in chromatin accessibility).

We also demonstrate that Foxa2 interacts with LXRα in ligand-dependent manner (**Fig. 3**). Our computational analysis provides evidence for 3 modes of synergistic regulation by Foxa2 and LXRα during ligand activation. Foxa2 initially bound DNA in complex with liganded LXRα, acting as a co-activator. LXRα either remains in the same position in the complex (**Fig. 5B**, top panel, IGV example on the right showing ligand-dependent Foxa2-only site that corresponds to increase in chromatin accessibility) or moves to an adjacent (**Fig. 5B**,, middle panel, IGV example demonstrating ligand-dependent dual Foxa2/LXRα binding site that corresponds to increase in chromatin accessibility) or a distal binding site (**Fig. 5B**, bottom panel, IGV example demonstrating ligand-dependent LXRα-only binding site that corresponds to increase in chromatin accessibility). A precedent for the first mechanism, showing that the forkhead consensus site but not the DR-4 LXR response element was required for T09-dependent induction of LPL promoter, with LXRα acting as a co-activator was reported by Kanaki and colleagues (*22*), We observe that forkhead motif is stronger in Foxa2/LXRα dual sites than nuclear receptor response elements. LXRα could be acting as a co-activator at some of those regions. In addition, we show a number of “Foxa2-only sites” containing strong forkhead motifs and lacking LXR response elements and LXRα signal. It is possible that the crosslinking is not sufficient to detect LXRα interacting with Foxa2 bound to DNA due to additional members of the complex. The evidence for the second mechanism is the presence of Foxa2/LXRα dual sites. Similarly, lack of co-localization, resulting in “Foxa2-only” and “LXR-only” could indicate a distal interaction where the proteins would be part of the same complex but their binding sites mapped to different genomic regions. Computational analysis that identified presence of binding sites of the same factors (from curated data) in “Foxa2-only” and “LXR-only” regions provides evidence for this possibility.

## Discussion

Our results challenge the classical paradigm of ligand-dependent activation of type II nuclear receptors, involving co-repressor/co-activator exchange, and initiation of transcription upon ligand binding (**Fig. 1A**). Unlike type I nuclear receptors, such as ER and AR, that change their subcellular localization and translocate to the nucleus upon ligand binding, LXR and other type II nuclear receptors including PPAR and FXR are always nuclear and, hence, should be able to access their binding sites. However, we demonstrate that extensive chromatin accessibility changes observed upon ligand activation are due to Foxa2 evicting nucleosomes to enable subsequent LXRα binding to DNA.

In addition, we determine a novel mechanism of nuclear receptor ligand-dependent binding. Foxa1, a close paralog of Foxa2. Foxa1 is required for binding of steroid receptors ER and AR and hormone-dependent gene activation by these receptors (*6, 23*). While Foxa1 is bound prior to ligand activation of steroid receptors and their translocation to the nucleus (*6, 7, 24, 25*), we find that Foxa2 occupancy is drastically induced by activation of type II receptor LXRα, suggesting there is an interdependent relationship between Foxa2 and nuclear receptor binding to DNA during ligand activation.

A recent report claimed that, like Foxa2, Foxa1 binding is hormone-dependent (*26*), but the data presented did not support those conclusions. First, the authors showed that an overwhelming majority of sites were bound by Foxa1 in both control and hormone treated conditions, while a small minority was condition-specific. Second, the minority of Foxa1 sites that were called ligand-dependent by ER and GR actually had read coverage in both basal and ligand-dependent conditions. The authors defined lack of binding by coverage below a certain cutoff rather than lack of ChIP coverage in unliganded state. Also, the authors did not provide Input controls to indicate what basal genomic coverage was expected. That study contradicted numerous previous reports showing that Foxa1 binding is ligand-independent) (*6, 7, 24*). In addition, a subsequent study resolved the controversy, showing that genome-wide Foxa1 binding is definitely hormone-independent and chromatin loops can create shadow Foxa1 binding sites (*25*). Hence, Foxa1 and Foxa2 indeed enable ligand-dependent binding of different types of nuclear receptors in unique ways.

Furthermore, we show that Foxa2 interacts with LXRα in ligand-dependent manner. We provide evidence for three possibilities for mechanism of Foxa2/LXRα interaction (**Fig. 5B**). LXRα could play the role of a co-activator without binding DNA (**Fig. 5B**, top panel) or both Foxa2 and LXRα need to be bound either in a proximal interaction (**Fig. 5B**, middle panel) or distal interaction (**Fig. 5B**, bottom panel. Additional subsequent experiments, such as RIME (Rapid immunoprecipitation mass spectrometry of endogenous proteins) (*27*), will identify co-factors in Foxa2/LXRα complex and Foxa2 Hi-ChIP, an assay combining ChIP with chromatin conformation (*28*), will distinguish distal interactions between Foxa2 and LXRα.

Ultimately, we demonstrate that ligand-dependent activation of LXR gene expression requires Foxa2. We have previously shown that activation of FXR, another type II nuclear receptor, with cholic acid is Foxa2-dependent ((*13*). Since deletion of *Foxa2* in the liver leads to mild cholestasis (*20*), we placed Foxa2-deficient mice on a diet enriched with primary bile acid cholic acid. Cholic acid, an FXR agonist, induced expression of over 7,000 genes in wildtype mice and over 2,500 of them were downregulated in *Foxa2* mutants. Considering that ligand-responsive activation of gene expression by both FXR and LXRα is Foxa2-dependent, it is likely that Foxa2 is required to modulate chromatin accessibility in a common mechanism mediating ligand-activation of type II nuclear receptors.

## Acknowledgements

We thank S. Srabani for helpful comments and technical assistance. We thank S Anakk and I. Schulman for critical reading of the manuscript. I.M.B. was supported by National Diabetes and Digestive and Kidney Diseases Institute R01 award DK121059.

## Author Contributions

J.K. performed data analysis, X.W. performed experiments, A.J.P performed data analysis, C. W. performed experiments, I.M.B. developed the project, performed experiments, analyzed data, and wrote the draft of the manuscript.

## Materials and Methods

### Mice

*Foxa2* mutant mice (Foxa2^loxP/loxP^;Alfp.Cre) were kindly provided by Klaus Kaestner (University of Pennsylvania). The derivation of the Foxa2^loxP/loxP^;Alfp.Cre mouse model has been reported previously (*17*). LXRα null mice (LXRα^-/-^) (*29*)were kindly provided by Ira Schulman (University of Virginia). (*17*). Mice were genotyped by PCR of tail DNA as described (*17, 29*). Male mice 8-12 weeks of age were used for all studies. All animal work was approved by Animal Care and Use Committee at UVa (protocol number 4162–03–20).

### ATAC-Seq

Nuclei were isolated from frozen liver tissue (50 mg), homogenized in PBS with Complete Protease Inhibitor (PI, Roche) using a glass Dounce homogenizer. The homogenate was collected into 1.5 mL Eppendorf tube and washed with PBS+PI (2000g for 3 min at 4^0^C). The pellet was resuspended in 1 mL Lysis buffer (10% NP-40, 10% Tween-20, 1% Digitonin) + PI and homogenized. The homogenate was transferred to 1.5 mL Eppendorf tube and placed on a rotator for 5min at 4^0^C. After centrifugation (2000g for 5 min at 4^0^C). the pellet was washed once with Resuspension buffer (1M Tris-HCl (pH 7.5), 5 M NaCl, 1 M MgCl2, 0.1% tween-20) and centrifuged (2000xg for 5min at 4^0^C). The resulting pellet was resuspended in PBS and filtered through cell strainer (40 μm). The nuclei were counted using Nexcelom cell counter. 50,000 nuclei were subsequently used in the tagmentation and library preparation protocol as described (*30*). Mitochondrial DNA contamination in amplified libraries was checked by QPCR before sequencing. All samples were sequenced on Illumina NextSeq 500.

### Chromatin Immunoprecipitation (ChIP) and ChIP-Seq

Snap-frozen mouse liver (100 mg) from wildtype and *Foxa2* mutant mice treated with vehicle and LXR ligands was used to prepare chromatin. ChIP and ChIP-Seq were performed as described previously (*31, 32*). Foxa2-specific rabbit antiserum (Seven Hills Bioreagents, WRAB-1,200) and rabbit polyclonal antibody specific to LXRα (Active Motif, 61175) were used for immunoprecipitation. All samples were sequenced on Illumina NextSeq 500.

### Coimmunoprecipitation (Co-IP) and Western Blotting (WB)

Mouse liver tissue was washed with cold PBS, and then lysed by homogenizing in cell lysis buffer (Cell Signaling Technology). After centrifugation at 14,000 rpm for 15 minutes at 4°C, the supernatants were collected, and the protein concentrations were measured using BCA protein assay reagents (Thermo Fisher). Subsequently, Co-IP was performed according to the immunoprecipitation protocol (Thermo Fisher) (thermo_sher.com/immunoprecipitation). Briefly, Antibodies were added to the protein G Dynabeads, incubated with rotation for 10 minutes at room temperature for binding. The beads-antibody complex was then crosslinked using the crosslinking reagent BS3 at room temperature for 30 minutes. After washing to remove non-crosslinked antibody, the antibody-crosslinked beads were incubated with equal amounts of protein lysates for overnight at cool room. The beads were washed for 5 time with 500 μL of IP cell lysis buffer and then eluted using 50 mM glycine. The eluted proteins were separated in Bolt 4–12% Bis-Tris gradient gel (Invitrogen), and transferred onto PVDF membranes (Azure). After blocking with 5% milk, the membranes were then probed at 4°C overnight with various primary antibodies: anti-foxa2 (Seven Hills) or LXR-α (Abcam), washed with TBST (20 mM Tris, 150 mM NaCl, 0.1% Tween 20; pH 7.6), and incubated with horseradish peroxidase (HRP)-conjugated secondary antibodies (Promega) at room temperature for 1 hr. Finally, after washing with TBST, the antibody-bound membranes were treated with enhanced chemiluminescent Western blot detection reagents (Azure) and imaged with Azure C300 Digital Imager.

### RNA Isolation and Sequencing

Liver RNA was isolated from Foxa2^loxP/loxP^;Alfp.Cre mice and control litter-mates as described previously (*17*). Quality of RNA samples was analyzed using Agilent RNA 6000 Nano Kit (Bioanalyzer, Agilent Technologies). Samples with RIN scores above 9.5 were used in library preparation. 1 μg of total RNA was used to isolate mRNA (NebNext Poly(A) mRNA Magnetic Isolation Module). Libraries of resulting mRNA were prepared using NebNext Ultra II RNA library preparation kit. All samples were sequenced on Illumina NextSeq 500.

### ATAC-Seq Analysis

Paired-end reads were aligned to the mouse genome (mm10; NCBI Build) using BWA (*33*). Duplicate reads were removed. Reads (phred score > 30) that aligned uniquely were used for subsequent analysis. Data from two biological replicates were merged for each condition (WT-Veh, WT-GW, WT-T09, Foxa2-KO-Veh, Foxa2-KO-GW, Foxa2-KO-T09). PeakSeq (*34*) was used to identify bound regions in ligand-activated condition against vehicle controls (WT-GW vs. WT-Veh & WT-T09 vs. WT-Veh, FDR 5%, q-value = 0.05).

### ChIP-Seq Analysis

Reads were aligned to the mouse genome (mm10; NCBI Build) using BWA (*33*). Duplicate reads were removed. Reads (phred score > 30) that aligned uniquely were used for subsequent analysis. For Foxa2 ChIPs, data from three biological replicates were merged for each condition (WT-Veh, WT-GW, WT-T09, LXRa-KO-Veh, LXRa-KO-GW, LXRa-KO-T09). For LXRa ChIPs, data from four biological replicates were merged for each condition (WT-Veh, WT-GW, WT-T09, Foxa2-KO-Veh, Foxa2-KO-GW, Foxa2-KO-T09). PeakSeq (*34*)was used to identify bound peaks against input controls (Foxa2 FDR 5%, q-value = 0.0005; LXRa FDR 5%, q-value = 0.005).

### RNA-Seq Analysis

RNA-Seq reads were aligned using STAR (*35*) to mouse genome build mm10. Expression levels were calculated using RSEM (*36*). Differential expression analysis of RNA-seq (*p*-value <0.05) was performed in R using EdgeR package (*37*) with a Benjamini-Hochberg FDR of 5%.

### Functional Analysis

Ingenuity curate’s information gathered from the literature, as well as genomic experiments (microarray, RNA-Seq, ChIP-Seq) to determine sets of targets controlled by a regulator (a transcription factor, chromatin remodeler, kinase, small molecule, etc.) and the logic of this regulation (activation or repression). IPA’s analysis method compares the overlap of each such set of targets with the subset of genes that are controlled by each regulator in an experimental gene list and reports the p-value of the overlap (Fisher’s exact test), If the overlap is significant, and the expression changes agree with those expected from the regulatory connection (genes activated by the regulator are activated in the analysis data set and repressed genes are repressed, for example), IPA analysis predicts whether the regulator associated with the gene targets is itself activated or inhibited. Analysis of overrepresented functional categories and upstream regulators in Ingenuity Pathway Analysis and heat map generation for RNA-Seq data was performed as described (*14*). Heatmaps of ChIP-Seq coverage were generated by deeptools (*38*). Chromatin-x enrichment analysis and analyses for over-represented pathways and disease conditions were performed by Enrichr (*15*). Sequence analysis for over-represented transcription factor-binding motifs in regions from ChIP-Seq experiments were performed by PscanChIP (*16*).

