## Supplementary material for "Pioneer factor Foxa2 enables ligand-dependent activation of LXRα": Figure S2

A

### Co-localized l Foxa2/LXR sites

First NR motif → HNF4G

| Name | ID | L.PV ▾ | L.O/U | G.PV | G.O/U |
|---|---|---|---|---|---|
| Foxa2 | MA0047.2 | 4.6E-208 | ↑ | 0 | ↑ |
| FOXA1 | MA0148.3 | 6.6E-203 | ↑ | 0 | ↑ |
| FOXK1 | MA0852.2 | 4.6E-164 | ↑ | 0 | ↑ |
| FOXO4 | MA0848.1 | 7.6E-157 | ↑ | 0 | ↑ |
| FOXO6 | MA0849.1 | 3.7E-154 | ↑ | 0 | ↑ |
| FOXI1 | MA0042.2 | 1.1E-152 | ↑ | 0 | ↑ |
| FOXP1 | MA0481.2 | 6.5E-144 | ↑ | 0 | ↑ |
| FOXL1 | MA0033.2 | 2.9E-135 | ↑ | 0 | ↑ |
| FOXB1 | MA0845.1 | 3.0E-134 | ↑ | 0 | ↑ |
| FOXD2 | MA0847.1 | 5.8E-133 | ↑ | 0 | ↑ |
| FOXK2 | MA1103.1 | 3.3E-132 | ↑ | 0 | ↑ |
| FOXD1 | MA0031.1 | 2.4E-131 | ↑ | 0 | ↑ |
| FOXC1 | MA0032.2 | 1.5E-125 | ↑ | 0 | ↑ |
| Foxj2 | MA0614.1 | 7.4E-124 | ↑ | 0 | ↑ |
| FOXC2 | MA0846.1 | 4.9E-115 | ↑ | 0 | ↑ |
| FOXO3 | MA0157.2 | 1.8E-114 | ↑ | 0 | ↑ |
| Foxo1 | MA0480.1 | 1.1E-102 | ↑ | 0 | ↑ |
| FOXP2 | MA0593.1 | 1.4E-100 | ↑ | 0 | ↑ |
| Foxj3 | MA0851.1 | 3.3E-100 | ↑ | 0 | ↑ |
| FOXP3 | MA0850.1 | 9.1E-93 | ↑ | 0 | ↑ |
| FOXF2 | MA0030.1 | 2.9E-90 | ↑ | 0 | ↑ |
| FOXG1 | MA0613.1 | 2.5E-84 | ↑ | 0 | ↑ |
| HNF4G | MA0484.1 | 8.5E-31 | ↑ | 2.3E-172 | ↑ |

B

### Foxa2 sites, no LXR signal

| Name | ID | L.PV | L.O/U | G.PV ▾ | G.O/U |
|---|---|---|---|---|---|
| FOXC2 | MA0846.1 | 2.7E-28 | ↑ | 1.4E-255 | ↑ |
| FOXC1 | MA0032.2 | 3.0E-29 | ↑ | 4.7E-255 | ↑ |
| FOXB1 | MA0845.1 | 1.7E-30 | ↑ | 6.4E-251 | ↑ |
| FOXA1 | MA0148.3 | 5.7E-44 | ↑ | 9.3E-230 | ↑ |
| Foxj3 | MA0851.1 | 3.8E-21 | ↑ | 2.4E-221 | ↑ |
| Foxa2 | MA0047.2 | 1.1E-44 | ↑ | 2.7E-220 | ↑ |
| FOXD2 | MA0847.1 | 7.7E-33 | ↑ | 1.9E-219 | ↑ |
| FOXL1 | MA0033.2 | 4.1E-33 | ↑ | 2.9E-214 | ↑ |
| FOXK2 | MA1103.1 | 1.1E-29 | ↑ | 2.7E-199 | ↑ |
| Foxj2 | MA0614.1 | 7.5E-24 | ↑ | 8.9E-192 | ↑ |
| FOXO4 | MA0848.1 | 2.7E-30 | ↑ | 4.7E-183 | ↑ |
| FOXI1 | MA0042.2 | 3.1E-32 | ↑ | 1.9E-175 | ↑ |
| Foxq1 | MA0040.1 | 2.3E-9 | ↑ | 3.0E-173 | ↑ |
| Foxd3 | MA0041.1 | 0.0002 | ↑ | 1.2E-171 | ↑ |
| FOXG1 | MA0613.1 | 6.3E-22 | ↑ | 2.3E-171 | ↑ |
| FOXO6 | MA0849.1 | 8.2E-30 | ↑ | 1.2E-170 | ↑ |
| FOXF2 | MA0030.1 | 5.1E-21 | ↑ | 1.9E-169 | ↑ |

**Figure S2**
