## Supplementary figures and images for "Pioneer factor Foxa2 enables ligand-dependent activation of LXRα"

### Figure S1

Foxa2 ChIP in LXRα KO

LXRα ChIP in Foxa2 KO

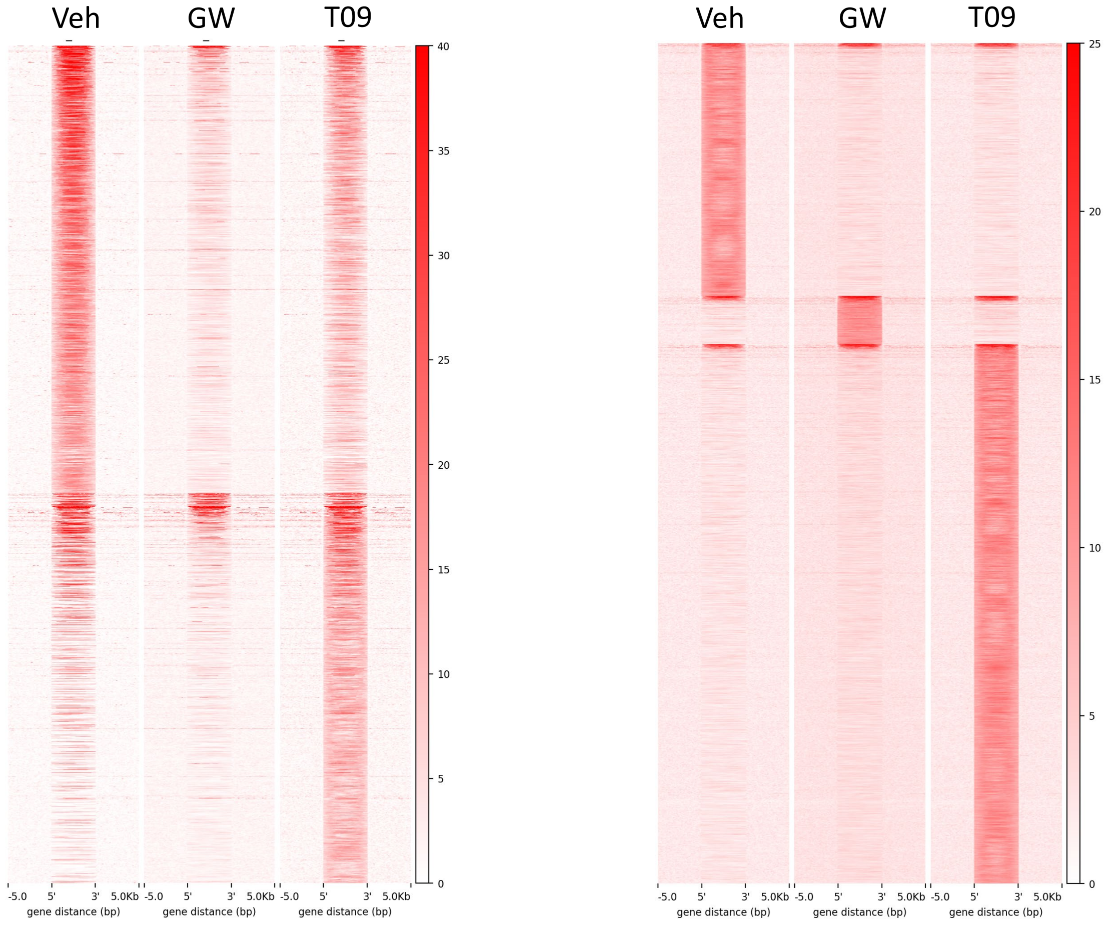


FDR 5%, q-value = 5E-4;

FDR 5%, q-value = 5E-3

**Figure S1**
